# Acute inhibition of the STA3 signaling pathway during epileptogenesis prevent GABAergic cells’ loss and imprinting of epileptic state: an in-vitro proof

**DOI:** 10.1101/2022.06.22.497204

**Authors:** S. Martín-Suárez, JM Cortes, P. Bonifazi

## Abstract

Epilepsy, the condition of recurrent unprovoked seizures resulting from a wide variety of causes, is one of the world’s most prominent brain syndrome. Seizures which are an expression of neuronal network dysfunction occur in a positive feedback loop of concomitant factors where seizures generate more seizures, including also neuro-inflammatory responses. Among other pathways involved in inflammatory responses, the JAK/STAT signaling pathway has been proposed to prevent epilepsy. In this work we tested on a model of temporal lobe epilepsy in-vitro, the hypothesis that acute inhibition of STAT3-phosphorylation - during epileptogenesis, can prevent structural damages in the hippocampal circuits, and the imprinting both of neural epileptic activity and inflammatory glial states. We performed calcium imaging of spontaneous circuits’ dynamics in organotypic hippocampal slices previously exposed to hyper-excitable conditions through the blockage of GABAergic synaptic transmission. Epileptogenic conditions lead to imprinted epileptic dynamics in the circuits in terms of higher frequency of neuronal firing and circuits’ synchronizations, higher correlated activity in neuronal pairs and decreased complexity in synchronization patterns. Acute inhibition of the STAT3-phosphorylation during epileptogenesis, prevented the imprinting of epileptic activity patterns, general cell loss, GABAergic cells’ loss and the persistence of inflammatory reactive glial states. This work provides further evidence that inhibiting the STAT3 signaling pathway under epileptogenesis can prevent patho-topological reorganization of neuro-glial circuits.

## INTRODUCTION

Epilepsy is a syndrome characterized by chronic aberrant patterns of cerebral activity, i.e. seizures, which display under variegate symptoms such as convulsions, loss of consciousness, mental absence and others. Typically, once appeared, epileptic patterns remain imprinted also under latent conditions in the brain, and re-display chronically possibly developing and can scaling up symptoms fatherly with additional dysfunctional problems (Covolan and Mello 2000; Curia et al. 2008; Fujikawa 1996; Lévesque, Avoli, y Bernard 2016; Lee et al. 2017). Epilepsy affects a large number of people estimated around over one percent according to a study of 2015 on the USA population (Zack and Kobau 2017) and represents the neurological disorder with largest numbers of patients. The relation between causes, prognosis and symptomatology is broad, complex and not fully understood (Stephen, Kwan, y Brodie 2001). Not all epileptic patients respond to the same way to drugs which should suppression or significantly reduce occurrence of seizures and repristinate life quality (Duncan 2006; Brodie et al. 2012). Anti-epileptic drugs can work through different mechanisms and often are aimed at preventing hyper-excitable brain conditions which are the fertile substrate for ictal patterns to appear and uncontrollably spread to the rest of the brain (Macdonald y Kelly 1995; Rogawski, Löscher, and Rho 2016).

Understanding the cause of epilepsy and the mechanisms underlying epileptogenesis, i.e. what turns a functional circuits into an epileptic one capable to generate ictal patterns of activity, is still a major open question which is the key for preventing and developing new effective treatments for epilepsy. Typical causes are related to head injuries, tumours, infections, stroke and genetics (Eyo, Murugan, y Wu 2017; Ahl et al. 2016).

The epileptogenic process leading to the final imprinting of an epileptic brain has been hypothesized to be a positive feedback loop (Fig. 1A) where seizures lead to more seizures through complex interactions which involve neuronal death, gliosis, emergence of aberrant connectivity and hyperexcitable conditions, neuroinflammation and other factors (Vezzani et al. 2011). In the case of temporal lobe epilepsy, one of the major forms of epilepsy, the circuit originating the seizures is located in the temporal region of the brain, specifically in the hippocampus. Although the causes which makes the hippocampus epileptogenic are not always necessarily known, and drugs are not always effective as treatment, its surgical removal can sometimes lead to the disappearance of the seizures, confirming its key role in generating ictal activity (Quirico-Santos et al. 2013)

**Figure 1.**
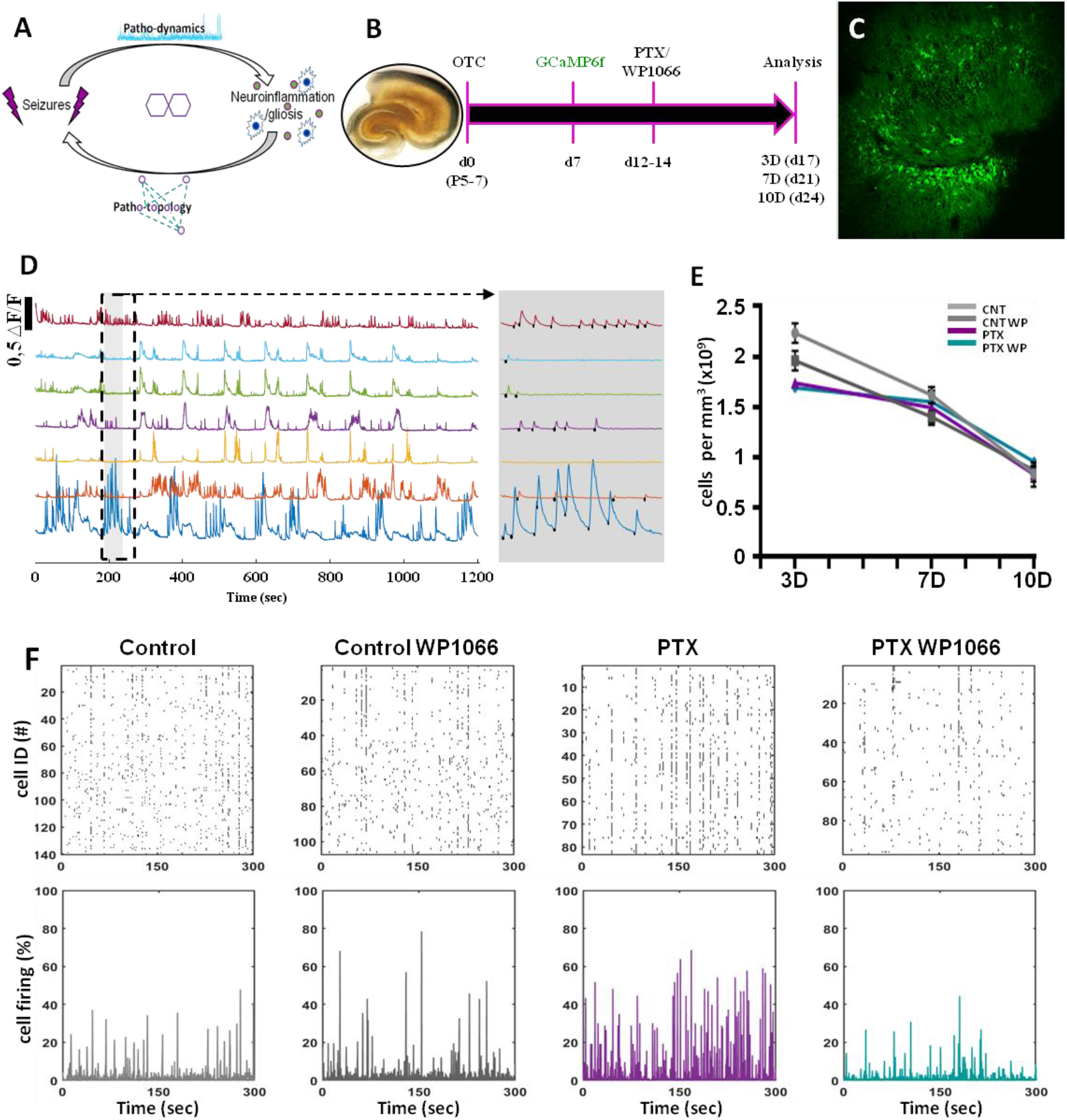
The experimental model: longitudinal calcium imaging and immunostaining. (A) Scheme of the positive feedback loop mechanism where seizures lead to more seizures through complex interactions which involve gliosis and neuroinflammation finally leading to patho-topological organization of neuro-glial circuit. (B) Scheme of the experimental procedure. The organotypic slices (OTC) extracted at P-5-7 (d0) and cultured. After 7 days (d7) slices are infected for the expression of the GCaMP6f. From d12 to d14 are exposed to PTX and/or WP1066, and imaging and immunostaining are performed 3-7-10 days after (i.e. at d17-21-24, we refer to them respectively as 3D, 7D and 10D). (C)Representative image acquired at 10x of the cells expressing the calcium sensor at d17. (D) Representative calcium traces of 7 neurons from a control slice at d17 with zoomed visualization (gray shaded) of the calcium traces and spikes’ onsets highlighted by black markers. (E) Health state of the circuits assessed by the density of alive cells per mm^3^. (F) Representative raster plots (top plots) showing the calcium spikes’ onsets of each cell over time in the four experimental group at d17. The percentage of active cell per frame is plotted in the bottom panels.

In this study we focused on possible mechanisms linking epileptogenesis to neuroinflammation mediated by the STA3 transcription factor, in the temporal lobe epilepsy context. Specifically, we used an in-vitro model of TLE, represented by organotypic hippocampal preparations exposed transitorily to hyper-excitable conditions (suppressing inhibitory synaptic communication). Hyper-excitable conditions are strongly epileptogenic and can trigger the emergence and imprinting of epileptic dynamics in circuits (Stoppini, Duport, y Corrèges 1997). In addition, organotypic hippocampal cultures have been considered too as in-vitro experimental models capable to generate per-se ictal-like patterns (Liu et al. 2017)

In our study, we focused on the impact of neuro-inflammatory response in the epileptogenic process when physiological circuits turn into pathological ones capable to display epileptic activity. In the last years clinical studies and experimental finding confirmed that the inflammatory pathways are implicated in the epilepsy. Several signaling pathways have been suggested to play an essential role in the development and progression of epilepsy and we focused on the JAK/STAT signaling pathway, in particular, on the STAT3 transcription factor. This pathway is involved in several process such as cellular proliferation, differentiation, neuron survival, death, and synaptic plasticity (Ghoreschi et al., 2009; Dedoni et al., 2010; Gupta et al., 2011; Makel a et al., 2010; Nicolas et al., 2012). In epilepsy STAT3 is highly expressed therefore its inhibition could reduce the neuronal damage and the progression of the epilepsy. The inhibitor of the STAT3 pathway, WP1066, has been reported to reduce the seizure severity and the progression of early epileptogenesis in a mouse model of epilepsy (Grabenstatter et al. 2014).

In our study, we hypothesized that the acute inhibition of Stat3 in presence of pro-epileptic hyper-excitable conditions, could attenuate the damage induced by the neuro-inflammatory response on the brain circuits preventing the manifestation of epileptic activity (Grabenstatter et al. 2014) and further damages such as cell loss.

Our results obtained imaging longitudinally the activity of circuits previously exposed to epileptogenic hyper-excitable conditions in simultaneous presence of STAT3 inhibitor, show that impacting the neuro-inflammatory response under epileptogenic conditions prevent the future appearance of hyper-synchronized patterns of activity, neural loss, inhibitory cells depletion and persistence of neuroinflammatory states. These results provide further evidence that neuroinflammation plays a key role on epileptogenesis and on the imprinting of epileptic dynamics, and that acute treatments impacting inflammatory responses can pre ent structural circuits’ damages and the appearance of dysfunctional activity patterns.

## METHODS

### HIPPOCAMPAL ORGANOTYPIC SLICE CULTURES

Slice cultures preparation was done according to what described in (Abiega et al., 2016). In brief, P5-7 Wild type pups were decapitated and the brains extracted and placed in cold dissetion medium (96% HBSS, 2% HEPES, 1% penicillin/streptomycin, 0,7% glucose (2,5M) and 0,3% of NaOH (0,5 M). Both hippocampi were dissected out and cut into 3 μm slices using a ibratome. Both hippocampi were cut in cold artificial aCSF (195mM Sucrose; 2.5 mM KCl; 1.25mM NaH_2_PO_4_; 28mM NaHCO_3_; 0.5mM CaCl_2_; 1 mM L-Ascorbic Acid/Na-Ascorbate; 3mM Pyruvic Acid/Na-Pyruvate; 7mM Glucose; 7mM MgCl_2_ in MiliQ), bubbled with 5% CO_2_ (Stoppini, Buchs, y Muller 1991). After slicing slices were then transferred to .4 μm culture plate inserts (Millipore, PICM01250). The membranes were placed in 24-well plates, each well containing 250μl of culture medium. The medium consisted of 50% Minimum Essential Medium supplemented with 2% B27, 25% horse serum, 2% Glutamax, 0,5% penicillin/streptomycin, 0.5% of NaHCO3, 0,5% glucose(2,5M), 0,8% sucrose (2.5M) and 18% HBSS. Slices were incubate at 37°C and 5% CO_2_ changing media the first day after doing the culture and every 2 days afterwards. Slices were kept in culture for 7 days before change to fresh culturing medium without B27 (Stoppini, Buchs, y Muller 1991; Cronberg et al. 2004). Epilepsy was induced by the addition of the GABA_A_ receptor inhibitor picrotoxin (PTX; 100 μM). PTX, vehicle or the inhibitor of Stat3 pathway, WP1066 (1,25 μM) was added with serum-free media during 3 days.

### CALCIUM IMAGING SET-UP AND RECORDING

For the calcium imaging, slices were infected with the calcium indicator AAV1.SYN.Gcamp6f.WPRE.SV40 ((Dana et al. 2019)) one week after the slicing. Twenty-four hours after the infection the medium was changed. Calcium imaging was performed using a 20x magnification objective on an inverted microscope (Zeiss Axio Observer.Z1 Apotome.2) for living cell imaging, equipped with a AxioCam MRc camera, a chamber for CO_2_ and temperature control for keeping same conditions as in cellular incubator. For imaging, the 24-well plate containing the slices from the same animal donor, and with covering the four different conditions studied in this paper (CNT, CNTWP, PTX, PTX-WP), was transferred to the microscope. The activity of each slice was then imaged for 20 minutes. One after the other all slices was imaged, and we did not follow any predefined sequence in the order of the slice to image. The plates were never opened, so the culturing conditions of the slices were always maintained to avoid any disturbance.

### CALCIUM IMAGING ANALYSIS

Neuronal cell body segmentation was performed as described previously in (Kanner et al. 2018). Briefly, the maximum of each pixel across all calcium images acquired for a gi en slice was used to reconstruct the image template and to segment neuronal cells’ contours through the custom-made software HIPPO. For each frame the average value of the pixels within a cell contour was calculated, and for each cell a calcium time series was then constructed across all frames. Time series were first high-pass filtered above 0.05 Hz to remove slow fluctuations and baseline changes, and next the traces were deconvolved using the ATLAB function “deconvolveCa” with default options, as derived and described from (Friedrich and Paninski 2016). The onsets of the calcium events were extracted from the deconvolved calcium signal with start and end points set by the respective threshold of 0.05 and 0.04 ΔF/F. The automatic detection of calcium spikes was later visually inspected for each cell. When the event detection was considered faulty, the starting and ending thresholds of a given calcium trace were adjusted manually always keeping respectively a 1 to 0.8 ratio. A binary time series representing the calcium activity in each frame was first reconstructed in each cell where the ones marked the onset of calcium spikes. Given a cell, the interval between two consecutive onsets was used as instantaneous firing rate (IF). In order to calculate the firing correlation in each neuronal pair, the binary time series were smoothed with a gaussian moving average using the MATLAB “smoothdata” function with a window length of 4 points and the correlation *C*_*ij*_ was calculated as

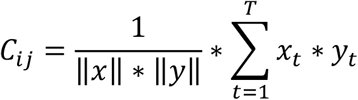

where *x*_*t*_ and *y*_*t*_ represent the time series of the neuronal pair *(i,j), T* the total number of frames (typically 4800 for a 20 minute recording) and the symbol represents the norm of a vector (i.e. the time series in the of *x*_*t*_ and *y*_*t*_).

In the case of global synchronizations, the binary time series were smoothed with a gaussian moving average using the MATLAB “smoothdata” function with a window length of 20, to link dynamics merging over larger time windows compared to neuronal pairs’ dynamics. As global synchronization index at time *t* (G*SI(t)*) we used the sum of the overall network activity obtained from the smoothed time-series (*s(t)*) according to:

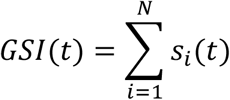

where *N* is the total number of imaged neurons and *s*_*i*_ is the smoothed time-serie of neuron *i*. Network synchronizations (*NSs*) were identified by *GSI* exceeding a threshold of chance level with p<0.05, as calculated from a thousand reshuffled network dynamics where single neuron time series were randomized while keeping the same inter-event distribution in each neuron.

To each *NS* was assigned the time frame of the peak of the corresponding *GSI*. All the cells recruited in a time window of seven frames around the *GSI* peak were considered as participating in the NS. The size of the NS was calculated as the percentage of cells participating in a given NS out of the total number *N* of imaged neurons in the circuit.

The frequencies of *NS* in a given circuit were calculated as the inverse of the intervals (in seconds) between consecutive *NSs*.

The similarity between two *NSs* was calculated as one minus the cosine between the binary vectors representing the cells participating in the synchronizations according to:

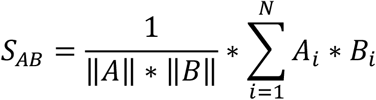

Where A and B represents the vectors of the two *NSs* and an element *A*_*i*_ (*B*_*i*_) is equal to one if the neuron *i* is participating in the network synchronization *A* (*B*).

The following table summarized the number of slices in each time point with the number of imaged neurons in each, and the relative experimental condition.

### INMUNOSTAININGS PROTOCOL

At the end of calcium imaging experiments, slices were washed with PBS and fixed with 4% paraformaldehyde for 30 min at room temperature. Immunohistochemical techniques were performed essentially as described before (Encinas and Enikolopov, 2008; Encinas et al., 2011). Slices were incubated with blocking and permeabilization solution containing 0.25% Triton-X100 and 3% bovine serum albumin (BSA) in PBS for 3 h at room temperature, and then incubated overnight with the primary antibodies (diluted in the same solution) at 4ºC. After washing with PBS, the sections were incubated with fluorochrome-conjugated secondary antibodies diluted in the permeabilization and blocking solution for 3h at room temperature. After washing with PBS, the sections were mounted on slides with Dako fluorescent mounting medium (Agilent-a o S3 3). The following primary antibodies were used: goat α-GFAP (Abcam Ab53554, 1:1,000); rabbit α-Iba 1 (Wako 19-19741, 1:1000); mouse α-NeuN (MerkMillipore, MAB377, 1:1000) Rabbit α-GABA (GeneTex GTX125988, 1:1000). The secondary antibodies used from ThermoFisher Scientific (1:1000), were: don ey α- rabbit Alexa Fluor 568 A 43) don ey α -mouse Alexa Fluor 647 (A-31571) goat α - rabbit Alexa Fluor 488 (A-34) and don ey α -goat Alexa Fluor 488 (A-84). 4′,6-diamidino-2-phenylindole (DAPI, Sigma-Aldrich D9542).

### IMAGE CAPTURE AND ANALYSI OF IMMUNOSTAININGS

All fluorescence images were collected employing a Leica Stellaris 5 (Leica, Wetzlar, Germany) microscope and LASAF software.

Images were exported as tiffs and adjusted for brightness and background, using Adobe Photoshop “le els”tool). All images shown are projections, from z-stacks of approximately 5 μm of thickness.

#### GABA quantification

Quantitative analysis of cell populations was performed by design-based (assumption free, unbiased) stereology using a modified optical fractionator-sampling scheme (previously described in (Encinas and Enikolopov 2008; Encinas et al. 2011).

#### Astrogliosis/microgliosis

The area occupied by astrocytes was measured in the slices using the open-source FIJI (Image J). GFAP+ -cells were selected in z-stacks using the “Threshold” tool to outline the pixels of the image labeled with GFAP. “easure” tool was used to calculate the % of occupied area by the staining in the image that have been highlighted before using the threshold (Martín-Suárez et al. 2020). Both measures were calculated from at least 5 z-stack from a minimum of three hippocampal slices.

#### Apoptosis

Apoptosis was quantified as previously described (Martín-Suárez et al. 2020). The number of pyknotic/ karyorrhectic (condensed/fragmented DNA) nuclei were quantified in the GCL+SGZ.

### STATISTICAL ANALYSIS

GraphPad Prism (Version 6 for Windows) was used for statistical analysis of the figure 5 and 6. One way ANOVA was performed in all cases to compare data from CNT, CNTWP, PTX and PTXWP in all the experiments. Error bars represent mean + standard error of the mean (SEM). Dots show individual data. To confirm the effect of the analysis Student’
ss t test was carried out. Results are expressed as means ± SEM. Significant data are denoted with asterisks: *P < 0.05; **P < 0.01; ***P < 0.001.

MATLAB was used for the statistical analysis of the following variables quantifying networ s’ dynamics: single neuron firing rate, neuronal pair correlation, network synchronizations (size and frequency), and synchronizations’ similarity.

The analysis was performed similarly in panel A of fig. 2, 3 and 5, and panel B and C of fig. 4, using pooled datasets. Specifically, given a variable, all the data obtained from different slices belonging to a given experimental group (CNT, CNT-WP, PTX, PTX-WP) and day of experiment (3D, 7D, 10D) were pooled. The statistical difference between groups was assessed using the Kruskall-Wallis’s test for non-parametric group comparisons with corresponding p-values for each pair of groups. Corresponding plots represent for each group medians, 25-75% percentile limits, smallest-highest values, and outliers.

**Figure 2.**
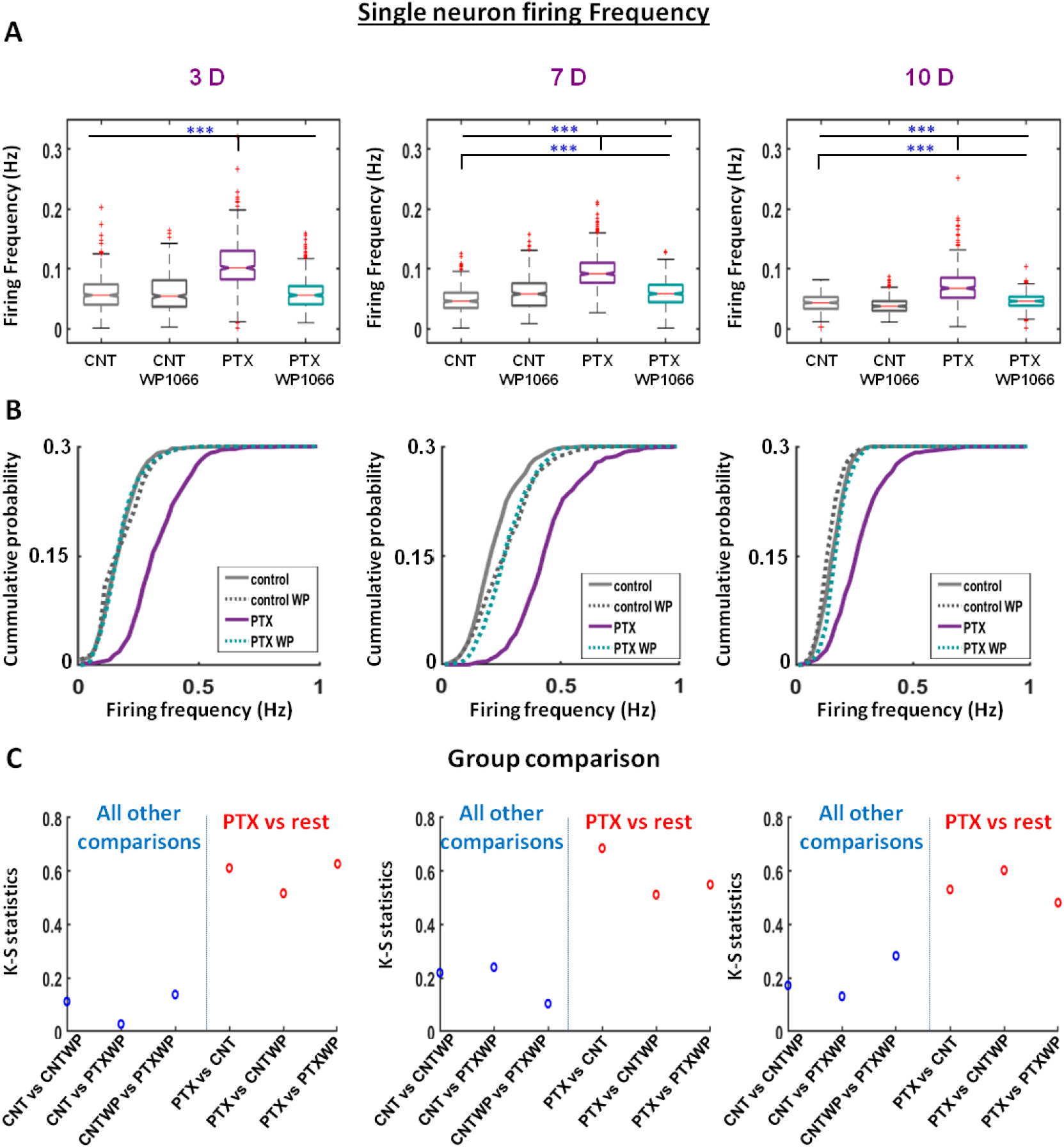
Single neuron level: frequency of calcium events. For each cell the average inverse of the intervals between consecutive calcium spikes are considered as firing rate. From left to right, results from slices respectively at 3, 7 and 10D are shown. (A) Pooled values at 3-7-10D across all slices are showing median (horizontal line) 25-75 percentile limits, bottom-top range values and outliers (marked by red asterisks). (B) Cumulative distributions obtained as average across slices belonging to same conditions. Fifty identical equally sized intervals were chosen within the minimum-maximum range across all groups. (C) Maximum difference between cumulative distributions shown in B (KS-S) for all group comparisons. Red dots highlight the comparison in the PTX-group vs all other groups. All other comparisons are shown as blue dots. Significant data are denoted with asterisks: *P < 0.05; **P < 0.01; ***P <0.001.

**Figure 3.**
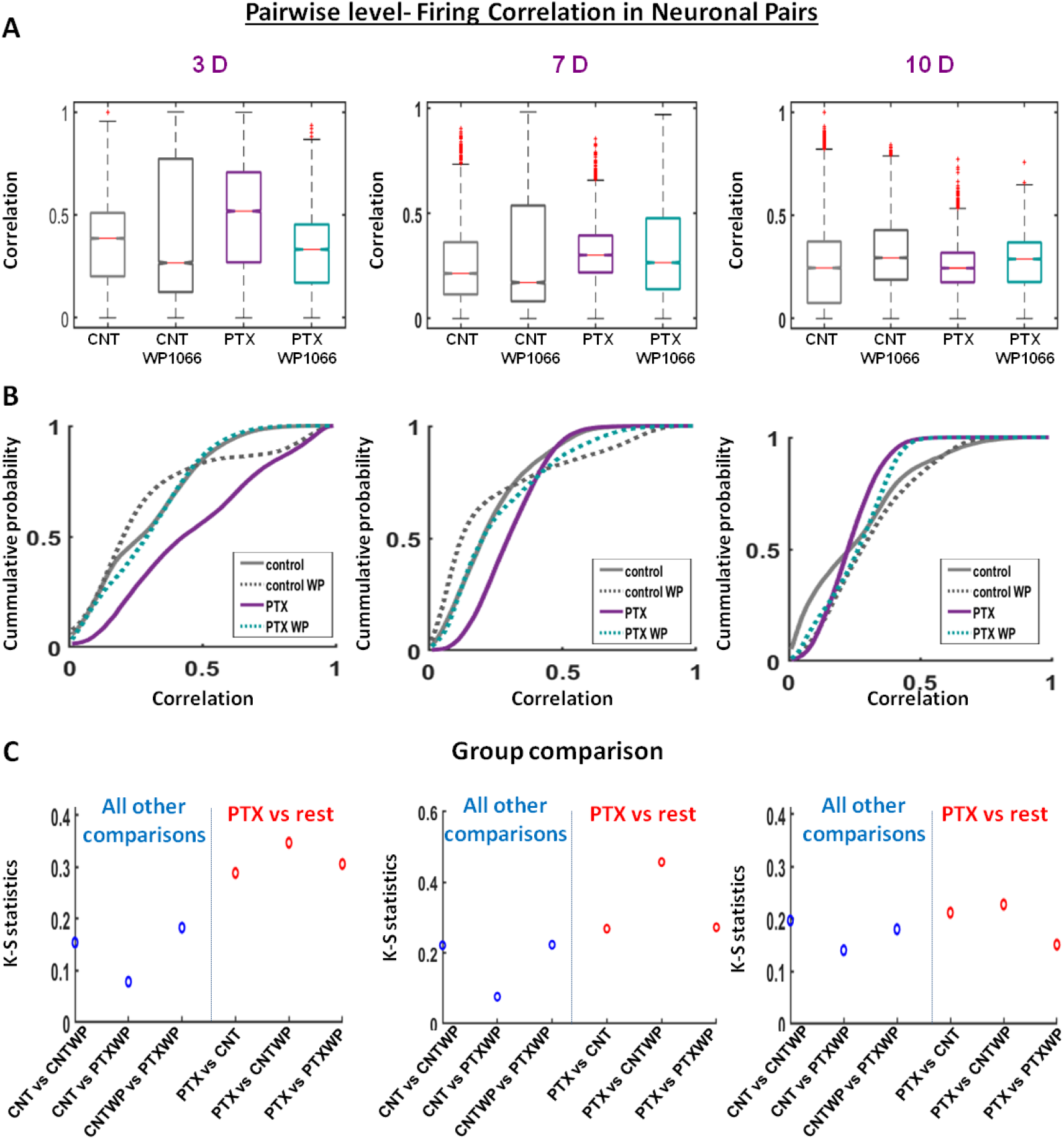
Pair-wise level: firing correlation in neuronal pairs’. Similar plot scheme to Fig.2. (A) Pooled values at 3-7-10D across all slices are showing median (horizontal line) 25-75 percentile limits, bottom-top range values and outliers (marked by red asterisks). (B) Cumulative distributions obtained as average across slices belonging to same conditions. Fifty identical equally sized intervals were chosen within the minimum-maximum range across all groups. (C) Maximum difference between cumulative distributions shown in B (KS-S) for all group comparisons. Red dots highlight the comparison in the PTX-group vs all other groups. All other comparisons are shown as blue dots. Significant data are denoted with asterisks: *P < 0.05; **P < 0.01; ***P < 0.001.

**Figure 4.**
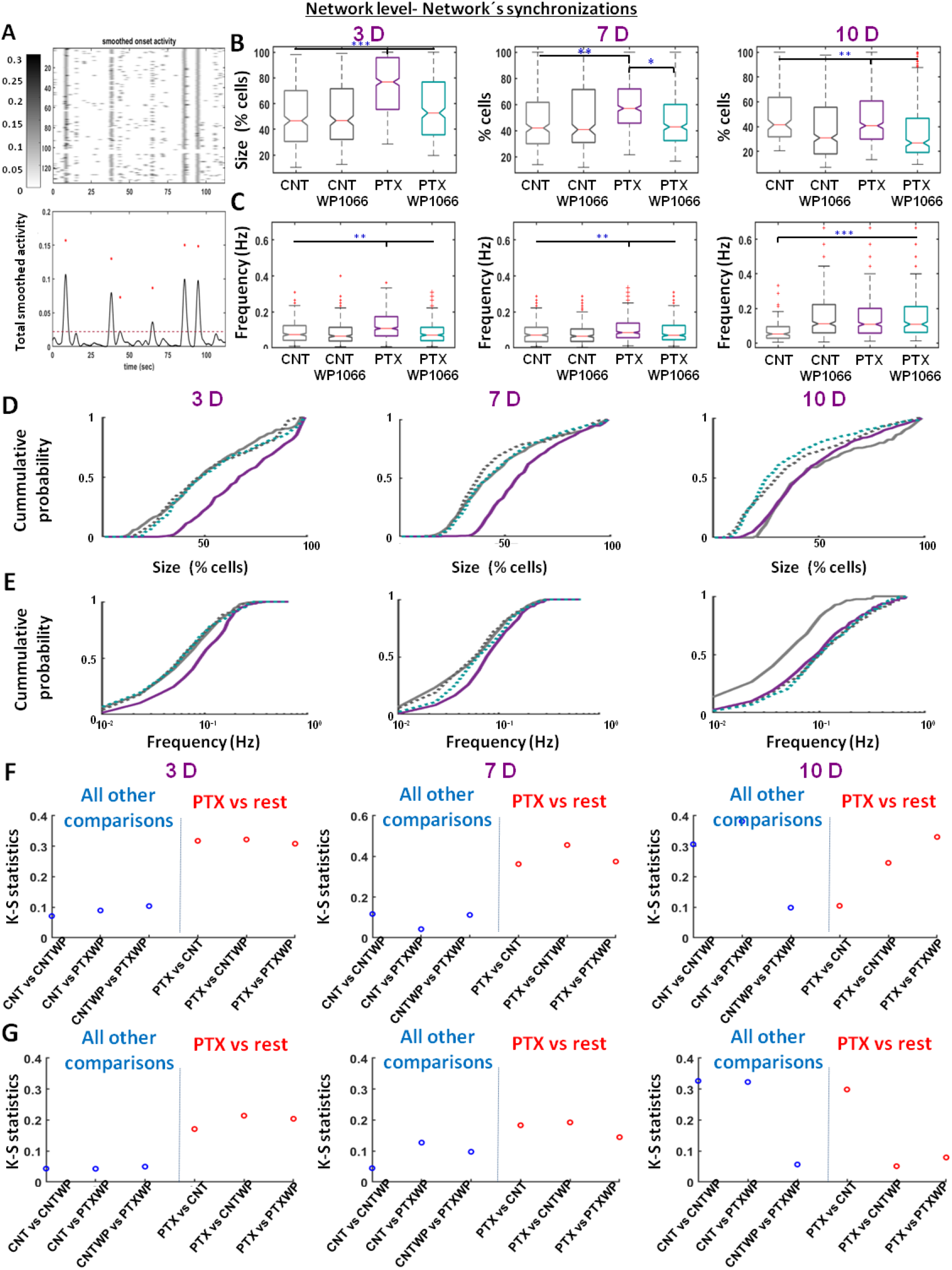
Ensemble level: size and frequency of circuits’ synchronizations. For panels B-G, a similar plot scheme to Fig.2 and 3 was used. (A) Representative zoomed raster plot (top) of the smoothed calcium spikes’ onsets and the sum per frame of the activity (bottom). The asterisks mark synchronization whose size are above chance level (marked as an horizontal line) (B, C) Pooled values of synchronizations’ sizes (B) and frequencies (C) at 3-7-10D across all slices are showing median (horizontal line) 25-75 percentile limits, bottom-top range values and outliers (marked by red asterisks). (D, E) Cumulative distributions of synchronizations’ sizes (D) and frequencies (E) obtained as average across slices belonging to same conditions. Fifty identical equally sized intervals were chosen within the minimum-maximum range across all groups. (F, G) Maximum differences (KS-S) between cumulative distributions shown in D (panel F) and E (panel G) for all group comparisons. Red dots highlight the comparison in the PTX-group vs all other groups. All other comparisons are shown as blue dots. Significant data are denoted with asterisks: *P < 0.05; **P < 0.01; ***P < 0.001.

Panel B of fig 2, 3 and 5, and panel D and E of fig. 4 represent the cumulative distributions obtained as average across slices for a given experimental group and day of recording. Specifically, the range of the considered variable (from the smallest to the highest value) was equally divided into fifty intervals. For each slice the cumulative distribution of the variable was calculated on such intervals. The average cumulative distribution across slices was then calculated and plotted.

**Figure 5.**
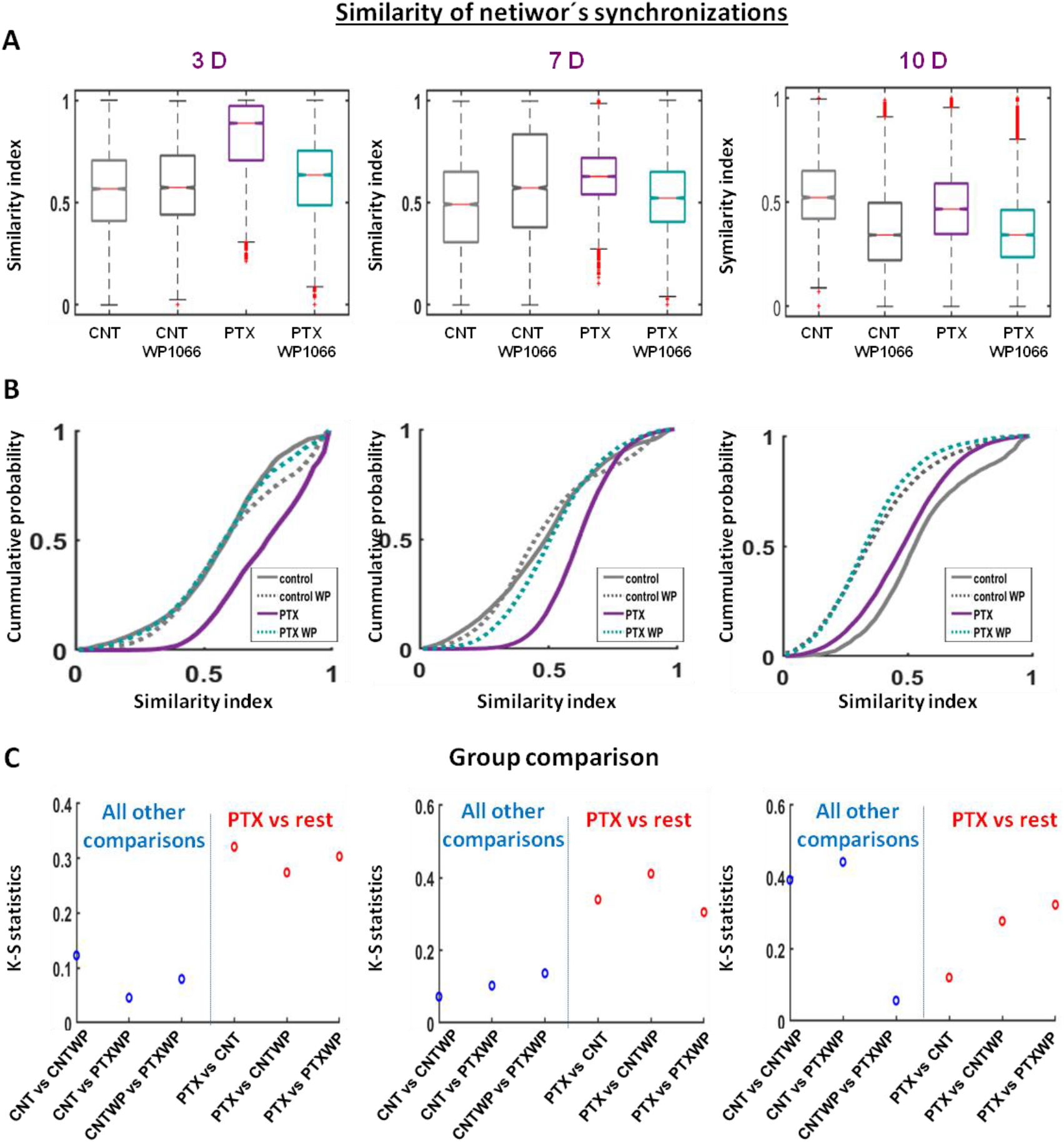
Ensemble level: similarity between circuits’ synchronizations. Similar plot scheme to Fig.2 and 3. (A) Pooled values at 3-7-10D across all slices are showing median (horizontal line) 25-75 percentile limits, bottom-top range values and outliers (marked by red asterisks). (B) Cumulative distributions obtained as average across slices belonging to same conditions. Fifty identical equally sized intervals were chosen within the minimum-maximum range across all groups. (C) Maximum difference between cumulative distributions shown in B (KS-S) for all group comparisons. Red dots highlight the comparison in the PTX-group vs all other groups. All other comparisons are shown as blue dots.

**Figure 6.**
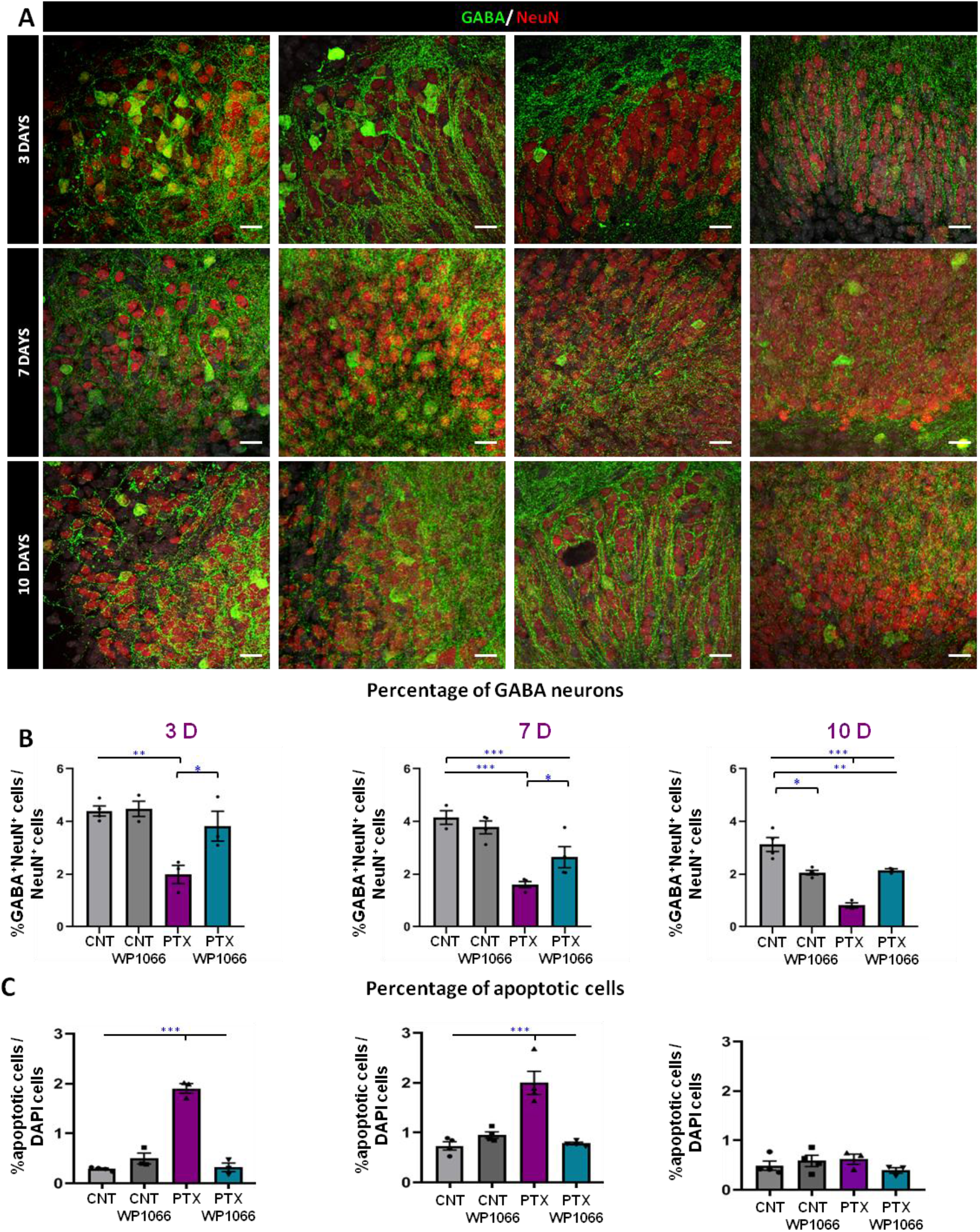
Characterization of circuits’ composition through GABAergic and apoptotic cell density: WP1066 prevents cell loss and death of GABAergic neurons. (A) Representative confocal microscopy images showing GABA positive cells in the granule cell layer in the different conditions at 3, 7 and 10 days. (B) Quantification of the proportion of GABA/NeuN positive cells among the total number of NeuN positive cells at 3, 7 and 10 days after the addition of PTX/WP1066. (C) Quantification of the proportion of apoptotic cells in the slices at 3, 7 and 10 days after the addition of PTX/WP1066. Scale bar is 10 μm. One-way ANOVA after all pairwise multiple comparisons by Holm-Sidak post hoc test. Bars show mean ± SEM. Dots show individual data. *p < 0.05, **p < 0.01, ***p < 0.001.

Panel C of fig. 2, 3 and 5 and panel F and G of fig. 4 plot the Kolmogorov-Smirnov statistics (KS-S) as a measure of distance between two experimental groups. Given a pair of experimental groups (out of CNT, CNT-WP, PTX and PTX-WP) the KS-S was calculated as the maximum difference between the corresponding cumulative distributions on each interval of the variable considered (Smirnov 1939).

## RESULTS

### EXPERIMENTAL MODEL: LONGITUDINAL STUDY OF CIRCUITS EXPOSED TO EPILEPTOGENIC CONDITIONS AND INHIBITION OF STAT3-PATHWAY

In order to assess the impact of inhibiting STAT3 signaling pathway during time windows of epileptogenesis, we studied in-vitro hippocampal organotypic slices extracted at P5-7 and cultured for at least 17 days (see Fig. 1A, B).

As model of epileptogenesis we exposed transitorily the circuits to hyper-excitable conditions, by blocking GABAergic transmission through Picrotoxin (an antagonist of the GABA-A receptor). Inhibition of the STAT3-pathway was performed using WP-1066, which inhibits the transcription of STAT3 gene (we also refer to it in the following as inhibited STAT3 condition, ISC, (Grabenstatter et al. 2014). To have full monitoring of all experimental conditions including needed controls, we designed accordingly four groups of experiments: a) control conditions (CNT group), where cultured slices were never exposed to any epileptogenic or inhibited STAT3-conditions; b) transitory epileptogenic conditions (EC; PTX group), where cultured slices were undergoing episodic suppression of GABAergic transmission in extracellular presence of PTX (100 μM) from DIV12 to DIV14) transitory epileptogenic and inhibited STAT3 conditions (EC-ISC; PTX-WP group), where slices were undergoing episodic suppression of GABAergic transmission as in the PTX group with the simultaneous extracellular application of the stat3 inhibitor WP-1066 (1,25 μM) and d) control under inhibited STAT3conditions (CNT-ISC; CNT-WP group), where slices were transitorily exposed to WP1066 (1,25 μM) from DIV12 to DIV14. In our protocol, the slices obtained from the same animal (batch) were separated to cover the four different experimental conditions, cultured on the same multi-well plate, and imaged on the same DIV (see Methods).

To monitor longitudinally the activity of epileptic circuits and the inhibition of the STAT3 transcription during hyper-excitable epileptogenic conditions, we performed calcium imaging (using GCaMP6f expressed only in neuronal cells; see Fig. 1B and Methods for viral infection) around the granular cell layer region (Fig. 1C) at 3, 7 and 10 days following conditioning with PTX/WP1066 (i.e. 17, 21 and 24 DIV, we refer to them as 3D, 7D and 10D). Spontaneous circuit’s dynamics were monitored in several dozens of neurons simultaneously over twenty minutes while fully preserving the culturing conditions (see Methods). Following semi-automated cell contour segmentation and calcium spikes’ detection (see Methods), the onsets of calcium events were extracted from each neuronal trace in order to reconstruct circuits’ dynamics from single neuron firing properties, to pair and ensemble synchronizations (see Methods and Fig. 1D and F). In addition, at 3D, 7D and 10D slices were also stained for immunochemical characterizations of inflammatory conditions in the glial cellular population and of circuit’s cellular state/composition (i.e. counting alive/apoptotic cells and GABAergic cells density; see Figure 1E, 6 and 7). Before describing the results of the study, Fig. 1E show the trend of the density of alive cells at 3-7-10D (i.e. when all the circuits’ characterizations were done), as a general marker of the health state of the experimental model. Note the clear decrease of cell density with time in culture, more marked passing from 7D to 10D.

### INHIBITION OF STAT-3 PATHWAY UNDER EPILEPTOGENESIS: IMPACT ON CIRCUITS’ DYNAMICS

We imaged an average of 98+/-22 neurons across all experimental conditions as summarized in the table 1, on a total of N=60 slices.

**Table 1.** For each group and day of recording, the total number of slices imaged, the imaged neurons per slice (ImN) and the number of circuit’s synchronizations (CS) occurred in the twenty minutes of recordings are reported.

| Condition<br>Day | Control |  |  | Control WP |  |  | PTX |  |  | PTX WP |  |  |
| --- | --- | --- | --- | --- | --- | --- | --- | --- | --- | --- | --- | --- |
|  | Slices | ImN | CS | Slices | ImN | CS | Slices | ImN | CS | Slices | ImN | CS |
| 3 DAYS | Slice 1 | 70 | 31 | Slice 1 | 71 | 56 | Slice 1 | 83 | 94 | Slice 1 | 56 | 36 |
|  | Slice 2 | 77 | 80 | Slice 2 | 66 | 58 | Slice 2 | 105 | 55 | Slice 2 | 97 | 56 |
|  | Slice 3 | 163 | 75 | Slice 3 | 150 | 44 | Slice 3 | 84 | 67 | Slice 3 | 105 | 53 |
|  | Slice 4 | 140 | 72 | Slice 4 | 107 | 57 | Slice 4 | 113 | 94 | Slice 4 | 112 | 96 |
|  | Slice 5 | 135 | 54 | Slice 5 | 109 | 67 | Slice 5 | 59 | 61 | Slice 5 | 79 | 53 |
|  |  |  |  |  |  |  | Slice 6 | 95 | 168 |  |  |  |
| 7 DAYS | Slice 1 | 99 | 55 | Slice 1 | 72 | 41 | Slice 1 | 70 | 94 | Slice 1 | 139 | 87 |
|  | Slice 2 | 100 | 74 | Slice 2 | 124 | 80 | Slice 2 | 92 | 94 | Slice 2 | 90 | 77 |
|  | Slice 3 | 105 | 64 | Slice 3 | 84 | 54 | Slice 3 | 96 | 63 | Slice 3 | 98 | 74 |
|  | Slice 4 | 58 | 43 | Slice 4 | 103 | 39 | Slice 4 | 100 | 72 | Slice 4 | 102 | 57 |
|  | Slice 5 | 75 | 38 | Slice 5 | 84 | 53 | Slice 5 | 107 | 74 | Slice 5 | 101 | 51 |
| 10 DAYS | Slice 1 | 73 | 19 | Slice 1 | 113 | 73 | Slice 1 | 77 | 65 | Slice 1 | 101 | 88 |
|  | Slice 2 | 107 | 24 | Slice 2 | 128 | 102 | Slice 2 | 66 | 71 | Slice 2 | 105 | 90 |
|  | Slice 3 | 101 | 42 | Slice 3 | 94 | 102 | Slice 3 | 102 | 127 | Slice 3 | 119 | 97 |
|  | Slice 4 | 125 | 37 | Slice 4 | 88 | 48 | Slice 4 | 90 | 95 |  |  |  |
|  | Slice 5 | 88 | 46 | Slice 5 | 104 | 83 | Slice 5 | 97 | 64 |  |  |  |

We first focused on single neuron dynamics under the four different experimental conditions at 3D, 7D and 10D (Fig. 2). In each neuron, we calculated the instantaneous frequency of calcium events (*IF*, i.e. the inverse of the time interval between consecutive calcium spikes) as an indicator of neural firing frequency and general circuits’ excitability.

Pooling *IFs* across all neurons from all slices within the same group and day of recording (N>=325, see table 1), non-parametric group comparison (Kruskal-Wallis test, see Methods), revealed a significant difference at 3D, 7D and 10D between the PTX-group (we refer to it in the following as the epileptic group) and all other groups (Fig. 2A). Although differences in some cases could be detected also between other groups’ comparisons, the median IF of the PTX-group (Fig. 2A, red horizontal line in the bar plots) was always the highest. Further quantification of the statistical difference between the groups was performed using the distance between average cumulative distributions (in this case data were not pooled but averaged across slices, so a representative average cumulative distribution for each group and recording day was reconstructed; see Methods and Fig. 2B). We used as a metric of distance (i.e. difference) between groups; the Kolmogorv-Smirnov statistics (KS-S), i.e. the maximum difference between cumulative distributions (see Methods and Fig. 2C). The KS-S values between the epileptic group (PTX) and all other groups (CNT, PTX-WP and CNT-WP; see red dots in Fig. 2C) were always (i.e. at 3D, 7D and 10D) higher when compared to the distance between the other groups (blue dots).

Overall, higher levels of single neuron firing in the circuits (PTX group) exposed to transitory epileptogenic conditions confirm its hyper-excitable epileptic state, which is prevented by the acute inhibition of the STAT3-mediated neuro-inflammatory response (PTX-WP group), therefore inducing inhibited STAT3 conditions during possible epileptogenesis keep single neuron firing similar to baseline control conditions. Next, we looked at the coordinated firing in the circuits, starting from the neuronal pairs’ level, calculating correlation between the time series of neuronal firing (see Fig. 3 and Methods).

Non-parametric group comparison on pooled statistic of firing’s correlations across all neuronal pairs and slices showed significant differences between all groups at 3D, 7D and 10D (Fig 3A). Note that the statistic on neuronal pairs scales as the square of number of imaged neurons so the number of pooled observations in each group and recording day was very high (>17530 neuronal pairs) compared to single neuron statistics. At 3D the median of firing’s correlations in the PTX-group was remarkably higher (>33%) when compared to the other cases (0.52 in PTX, 0.39 in CNT, 0.23 CNT-WP and 0.34 in PTX-WP group respectively). Also, at 3D the KS-S revealed higher differences between the epileptic group compared to other groups as shown in Fig 3B and C. The same trend but attenuated was observed at 7D with higher pair correlation (>11%; 0.30 PTX, 0.22 CNT, 0.18 CNT-WP, 0,27 PTX-WP group) and higher KS-S when comparing the PTX-group compared to the others. At 10D both medians of IF and KS-S differences showed similar values when comparing groups. Overall, higher levels of pair-wise synchronized dynamics were observed in the epileptic group at 3D and 7D, but not at 10D (when also the density of alive cells in the circuits displayed lowest values, as previously shown in see Fig. 1E).

Next, we looked at larger ensembles’ dynamics and focused on circuits’ synchronizations (CS, Fig. 4). Since synchronized events could also occur by chance, we considered only synchronizations with sizes above chance level (Fig. 4A) as estimated from reshuffled random firing (see Methods). As first step, we quantified the frequency (panels 4B, D and F) and the size (panels 4C, E and G) of the synchronizations, the latter quantified as percentage of recruited cells within the imaged neural population.

The pooled statistics of circuits’ synchronizations showed similar trends at 3D and 7D, in fact non-parametric statistics showed that PTX-groups were significantly different to all other groups in terms of frequency and size of circuits synchronization (p<0.05) while all other groups did not show significant differences between them (apart CNT-AIC and PTX-AIC at 3D when p=0.05). Higher frequency of synchronizations (medians at 3D and 7D of PTX-group are 0.11 and 0.09 Hz respectively while all other groups are 0.07 Hz) and percentage of recruited neurons in synchronized events (medians of groups at 3D: PTX 76.8%, CNT 46.6%, CNT-WP 46.8%, PTX-WP 52.7%; medians of groups at 7D: PTX 57%, CNT 42.0%, CNT-WP 40.8%, PTX-WP 42.9%) were observed in the PTX-group compared to the other groups. Similarly, at 3D and 7D, the KS-S showed higher differences between the PTX-group and all other groups (Fig. 4B and C). At 10D, such trends were not observed anymore, and the pooled statistics revealed a significant difference between CNT and all other groups in terms of synchronizations frequency (Fig. 4C), while in terms of synchronization size all groups were significantly different apart the case of CNT vs PTX and CNT-WP vs PTX-WP (Fig. 4 D). Similarly, at 10D also the KS-S did not show the previously trends observed at 3D and 7D. Overall, and similarly to the correlated dynamics observed in neuronal pairs, synchronized dynamics of larger neural ensembles in the epileptic group were more frequent and recruited in a larger number of neurons at 3D and 7D, while at 10D such trends did not persist.

Finally, in order to characterize the complexity of the spontaneous ensembles’ dynamics in terms of richness of repertoire of patterns of synchronizations generated by the circuits, we looked at the similarity of the circuits’ synchronizations (Fig. 5), measured as the cosine of the angle between the binary vectors representing the neurons recruited in the circuits’ synchronizations (see Methods). The pooled statistics revealed significant differences between all groups at 3D, 7D and 10D (Fig. 5A). Note that also in this case as in the of neuronal pairs’ correlation, the number of observations (>3008 CS pairs) scales as the square of the total number of synchronizations. At 3D the similarity of the synchronizations generated in the PTX group was very high (median 0.89 out of a possible maximum of 1) compared to the other groups (0.57 CNT, 0.57 CNT-WP and 0.64 PTX-WP). The gap was attenuated at 7D (0.63 PTX, 0.49 CNT, 0.57 CNT-WP, 0.52 PTX-WP). The KS-S showed clearly how the distribution of events both at 3D and 7D in the PTX group was consistently more distant to all other groups (fig 5 B and C). All the above trends on the similarity of synchronizations patterns were not present at 10D.

### INHIBITION OF STAT-3 PATHWAY UNDER EPILEPTOGENESIS: IMPACT ON CIRCUITS’ STRUCTURE AND INFLAMMATORY STATE

Since the PTX-group displayed patterns of activity stereotypical of hyper-excitable epileptic states from single neuron firing, up to pair-wise correlations and larger ensembles’ synchronizations, we next performed immunostaining quantifications related to the circuits’ structure, specifically on the density of apoptotic cells and of the inhibitory neurons (i.e. GABAergic cells). Coherently with the reported literature (Ben-Ari y Represa 1990; Knopp et al. 2008; Houser 2014) on the GABAergic neurons’ loss across the development and installment of epilepsy, we hypothesized that: 1) PTX induce the loss of GABA cells leading to hyper-excitable circuits’ states, and that 2) inhibited STAT3 conditions during the time window of epileptogenesis reduce cellular damages (and the consequent impact on the emergence of epileptic circuits’ dynamics previously shown in Fig. 2-4).

Therefore, we quantified the number of GABA cells (identified by GABA staining, see Methods) in the granular cell layer, out of whole population of neurons (NeuN positive cells; Fig. 5A). At 3D we saw that the presence of GABA cells in the epileptic group was around 2%, about 50% less when compared to all other conditions where GABAergic population was about 4% of the overall neural population (Fig. 5B). Similar results additionally characterized by a slightly larger general decrease in the GABAergic population (more evident in the PTX-WP group with values around 3%) were obtained at 7D (central plot of Fig. 5B). Also, at 10D we observed under all conditions a general decrease of the percentage of GABAergic neurons, but still maintaining the trend of 3D and 7D with lowest values observed in the PTX-group (right plots of Fig. 5 B).

Since GABAergic cells presence was reduced under epileptic conditions but preserved by the acute inhibition of the STAT3, next we looked at overall cell loss, and we quantified the presence of dead cells at 3, 7 and 10D counting the number of apoptotic cells GCL+SGZ as a readout of cellular damage through the nuclear staining DAPI (Fig. 5C). We found that at 3D and 7D cell loss was higher in the epileptic group, with a higher significant ratio of apoptotic cells compared to the other conditions. This result highlights a general structural circuit’s damage in the epileptic group. The inhibiton of STAT3 under epileptogenesis (PTX-WP group) prevented general cellular damage. At 10D whole experimental group showed similar presence of dead cells, but this also corresponded to lowest and similar densities of alive cells in the cultures (as previously shown in Fig. 1E).

Overall, the above immuno-characterizations showed that epileptic conditions induce a general loss of cells targeting preferentially the GABAergic neuronal population which is decreased to half of the baseline levels, and that anti-inflammatory conditions prevent both overall cell loss and specifically of GABAergic cells.

Since we studied the impact on the epileptogenic conditions of inhibiting the STAT3-mediated neuro-inflammatory response, next we characterized the inflammatory profile of the circuit in the different glial populations. Reactive glia, involving both astrocytes and microglia, is one of the hallmarks of the epilepsy. We first focused on astrogliosis measuring the increased overall area of GFAP staining at 3D, 7D and 10D, which quantifies the level of reactive astrocytes (Fig. 7A). At 3D, 7D and 10D the expression of GFAP was significantly higher in the PTX group compared to all other groups, as measured by the area occupied by GFAP^+^ pixels (Fig. 7B and see Methods). However the inhibition of STAT3 reduces the area occupied by GFAP.

**Figure 7.**
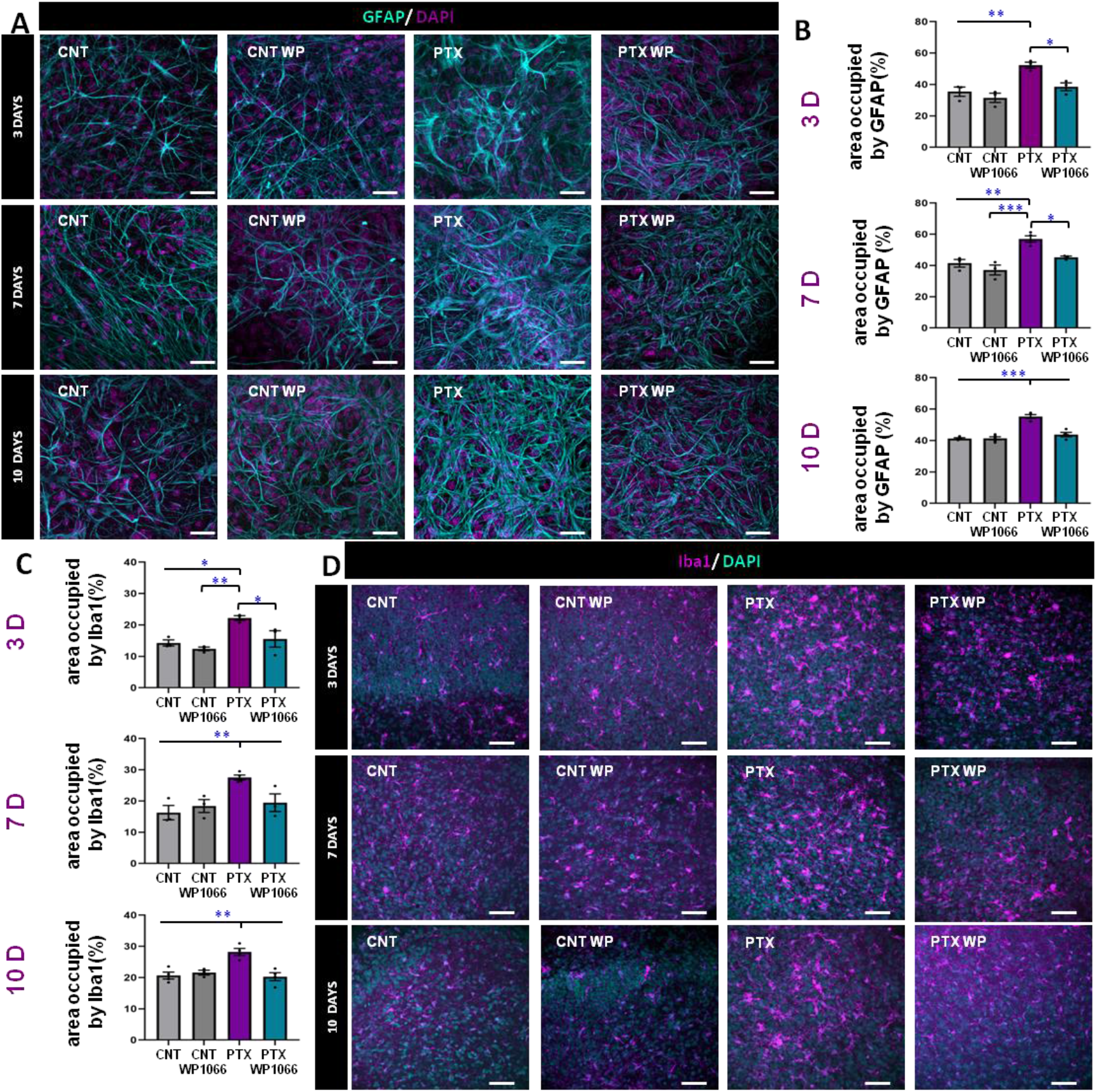
Circuits’ neuro-inflammatory profile: WP1066 prevent the glial inflammatory state which is present in epileptic circuits. (A) Representative confocal microscopy images showing the area occupied by the astrocytic marker GFAP in the different conditions at 3, 7 and 10 days. (B) Quantification of area occupied by GFAP at 3 days, 7 days and 10 days after the addition of PTX/WP1066. (C) Quantification of area occupied by Iba1 at 3 days, 7 days and 10 days after the addition of PTX/WP1066. (D) Representative confocal microscopy images showing the area occupied by the microglial marker Iba1 at 3, 7 and 10 days. (E) Confocal microscopy projections showing DAPI-stained condensed nuclei. (F) Quantification of apoptotic cells in the slices at 3, 7 and 10 days in the 4 different conditions. Scale bar is 10 μm. One-way ANOVA after all pairwise multiple comparisons by Holm-Sidak post hoc test. Bars show mean ± SEM. Dots show individual data*p < 0.05, **p < 0.01, ***p < 0.001.

Next, we tested the reactivity of the microglía cells, using the specific microglial marker Iba 1 (Fig. 7C). Similarly to the observation on the astrocytes’ reacti ity, we found that the expression of Iba1 as measured by the area occupied by the marker, is significantly increased in the PTX group compared to all other groups at 3D, 7D and 10D (Fig. 7D). The inhibition of STAT3 reduces the area occupied by Iba1.

The above overall results on glial reactivity show that the sole inhibition of the STAT3 transcription was capable to prevent the reactivity both in astrocytic and microglial cells under epileptogenic conditions.

The overall immunostaining characterizations of the structural and inflammatory profiles of the circuits show that hyper-excitable conditions induce a general neuroinflammatory profile and the gliosis is also accompanied by an increase in the number of apoptotic cells with higher loss in GABAergic population, which are stereotypic hallmarks of the epileptic state. Notably, we stress again that the sole inhibition of the STA3 transcription was capable to prevent the induction of these epileptic markers.

## DISCUSSION

We used organotypic hippocampal cultures as an in-vitro model of temporal lobe epilepsy, similarly to what previously described in literature (Stoppini, Buchs, y Muller 1991; Stoppini, Duport, y Corrèges 1997; Abiega et al. 2016). We exposed transitorily cultured circuits to hyper-excitable pro-epileptic conditions by suppressing GABAergic synaptic transmission to model epileptogenesis. Using calcium imaging to monitor spontaneous neural dynamics simultaneously on few dozens of neuron from 3 to 10 days following epileptogenetic episodes. We observed that epileptic circuits displayed increased single neuron firing, increased firing correlations in neural pairs, higher frequency of circuits’ synchronizations with higher number of recruited neurons and decreased complexity of synchronized patterns (i.e. a less rich repertoire of activity as reflected by higher similarity between generated patterns). In addition, in agreement with previous literature epileptic circuits displayed also higher cellular loss and in particular in regard to the GABAergic population(Knopp et al. 2008). Therefore, a lower presence of inhibitory cells was present in the circuits, favouring the appearance of hyper-excitable states with increased frequency of single cell and coordinate circuit’s activity. Moreover, epileptic circuits displayed a persistent inflammatory glial profile as revealed in astrocytic and microglia markers.

Although such observations in in-vitro hippocampal slices are in general agreement with previous literature (Duport, Stoppini, y Corrèges 1997; Stoppini, Duport, y Corrèges 1997) on in-vitro epileptic models, to the best of our knowledge this is the first time that spontaneous synchronizations and emergent collective dynamics were studied and quantified consistently in epileptic circuits longitudinally over days after epileptogenesis. This was possible by maintaining the culturing conditions during the imaging sessions, i.e. in un-perturbate conditions, avoiding perfusion of extracellular ACSF or medium which is the typical protocol used in previous works (Stoppini, Buchs, y Muller 1991)

When we blocked the transcription of the STAT3 gene, concomitantly during the time windows of epileptogenesis (by simultaneous application of PTX and WP1066), we observed that epileptic dynamics, cellular loss, GABAergic cells depletion, and neuro-inflammatory glial states were not present in the neuro-glial circuits, with ariables’ values at baseline control levels. This was clear longitudinally at 3 and 7 days following the episode of epileptogenesis while at 10 days no clear differences and trends could be anymore observed anymore. The loss of difference between the experimental conditions at 10D can be explained by the progressive cellular loss and decrease density of alive cells in the cultured circuits, which was occurring similarly in control baseline conditions and in all other tested conditions.

Organotipic hippocampal slices are commonly used as an in vitro model of seizures and chronic epilepsy (Abiega et al. 2016). Here we showed that organotypic cultures treated with PTX recapitulate the most typical features of the epiletogenesis: spontaneous electrical seizures, cell death, inflammation, and alteration of the neuronal circuits, as it was previously described by (Duport, Stoppini, y Corrèges 1997; Stoppini, Duport, y Corrèges 1997). Although there is some controversy about the intrinsic epileptic profile of organotypic slices even in control condition(Liu et al. 2017), our results demonstrated that organotypic slices treated with PTX is an excellent model to study and understand epileptogenesis.

Here we focused on the role of STAT3 in the development of epilepsy and neuroinflammation. It is known that JAK/STAT3 pathway is rapidly activated in the hippocampus after status epilepticus (Raible, Frey, and Brooks-Kayal 2014). Previous reports linked the activation of STAT3 to epileptogenesis (Okamoto et al. 2010)., Xu et al., suggested that STAT 3 remains activated up to 1 month which maintain and aggravate the effect pf astrogliosis (Xu et al. 2011). STAT3 can be inhibit pharmacologically by different drug such us Pyridone or WP1066. Grabenstatter et al. discovered that the inhibition of STAT by WP1066 lowered the number of spontaneuous seizures in a pilocarpine model of epilepsy in rats reducing the severity of the next seizures (Grabenstatter et al. 2014).

We herein describe that the inhibition of STAT3 with WP1066 not only reduced the effect of epilepsy on the neuronal circuits functionality but also we were able to maintain the topology of the circuit. We show that the alteration of the neuronal circuits is accompanied by topological changes such as the loss of GABAergic cells. We observed that the number of GABAergic cells decreased in comparison to control condition in all time points interestingly this loss of GABAergic cells has been shown in in vivo models of TLE induced by pilocarpine (Knopp et al. 2008). The role of GABAergic neurons in the development and prevalence of epilepsy has been widely studied in animal model, demonstrating that the loss of GABAergic interneurons may shift the balance between excitation and inhibition (Cohen et al. 2002; Ben-Ari and Represa 1990; Knopp et al. 2005). We demonstrated for the very first time that the inhibition of WP1066 reduced the loss of GABAergic cells in an in vitro model of epilepsy.

Neuroinflammation is another hallmark of epilepsy; therefore we resorted to analyze the inflammatory profile of the organotipic slices. Gliosis includes both, reactive astrocytes and reactive microglia since astrocytes and microglia are interconnected in the way that the activity of one type of cell affects the activity of the other (Sanz y Garcia-Gimeno 2020). We analized the area occupied by astrocytes and microglia as measurement of gliosis. Our results confirmed the results obtained previously by Xu et al., in which inhibition of STAT3 reduces the overexpression of GFAP (Xu et al. 2011) and Iba1 after epilepsy. Thus, we have demonstrated that the inhibition of STAT3 could impact in the activation of astrocites and microglia in the epileptogenic process.

Our data suggest that the inhibition of JAK/STAT at the onset of the epileptic event could modify the progress of epilepsy. The inhibition of this pathway may have a therapeutic value as complementary therapy to the commonly used anticonvulsant drugs.

